# Productivity drives demography and sex structure in a declining waterfowl species

**DOI:** 10.64898/2026.06.15.732236

**Authors:** Alain Caizergues, Alicia Lamé, Adrien Tableau, Guillaume Souchay

## Abstract

Demographic models are crucial for uncovering the mechanisms underlying changing populations trajectories and structure and to identify the drivers of such changes. Common pochard populations of North east/North west Europe have experienced a sharp decline accompanied by an increase in male proportions among adults since the mid-1990. We used a two-sex, two-stage deterministic matrix population model, to perform a prospective perturbation analysis and to explore, through simulations, some plausible causes underlying the observed decline and change in sex structure. We show that Common pochard populations are more sensitive to changing survival than to changes in productivity’s components (clutch size, nest survival…). However, due to an environmental variance much higher than that of survival, components of productivity, especially nest survival, would be the main drivers of Common pochard populations’ growth rate, a finding supported by empirical data. More importantly, we show that although sex-specific changes in survival at any stage of the life cycle are potent drivers of both population growth rate and changing sex ratio, there is no need to resort to them for explaining the increasing proportions of males such as observed in Common pochard. Because adult males display higher survival than adult females (0.74 against 0.64 on average), any factor affecting recruitment (nest or first year survival) increases the weight of adults into the populations and hence the proportion of males. Thus, in species displaying sex-biased mortality, such as many ducks, decreasing recruitment can underly declining population size and changes in sex structure at the same time, emphasising the importance of accounting for males in monitoring schemes and demographic models.

## Introduction

Biodiversity collapse worldwide is among the top most worrisome consequence of the impact of humans on the functioning of ecosystems and hence the future of the biosphere (Ceballos & Ehrlich, 2023). Over the years therefore, monitoring biodiversity has become a priority for: documenting the decline of species (Fujisaki et al., 2008, Collen et al., 2009, Folliot et al., 2022), understanding the mechanisms thereby populations collapse or resist anthropogenic pressures and to design mitigating actions (Elmberg et al., 2006). To achieve these goals, monitoring schemes have been implemented all-over the world. Many of these monitoring schemes simply consist in populations censuses (counts) coordinated over countries in order to cover whole populations. For example, the International Waterbirds Census (IWC) coordinated by Wetlands International covers more than 100 countries in Europe, Africa and Asia, while the Mid-Winter Waterfowl Survey (MWS), its North-American counterpart, is coordinated over all USA states. Such monitoring schemes proved invaluable for assessing the level of threat to species/populations in red lists (IUCN, 2023), and, to some extent, for evaluating the impact of climate change (Lehikoinen et al., 2013, Pavón-Jordán et al., 2019, Meehan et al., 2021), or measuring the effects of management actions (Gaget et al., 2020, Pavón-Jordan et al., 2020). However, counts provide limited insight into the underlying mechanisms of population changes, thereby impairing the understanding of populations’/species’ collapse/resilience. To properly understand demography, one must determine the relative influence of each demographic parameter such as survival, age at first breeding, fecundity… on population growth rate (***λ***), a procedure called prospective analysis (Caswell, 2019). By allowing to rank the effects of each demographic parameters on ***λ***, prospective analysis helps assessing the anthropogenic or natural drivers of population size and hence to design suitable management actions (e.g., Flint et al. 2015). Species can be ranked along a slow-fast demographic continuum that depends on their generation time and explains their relative sensitivity to changes in productivity *versus* changes in adult survival (Lebreton & Clobert, 1991, Koons et al., 2014). The fitness of species displaying a generation time below 2 years depends more on changing productivity than on changing adult survival; thus, they tend to buffer productivity from environmental stochasticity (Gaillard et al., 2000, Koons et al., 2014, Hilde et al., 2020). Conversely, species whose generation time is above 2 years are more sensitive to changing adult survival and tend to buffer it from environmental stochasticity (Gaillard et al., 2000, Koons et al., 2014, Hilde et al., 2020).

Although being powerful tools for assessing demographic drivers, prospective analyses often suffer shortcomings. Firstly, males may be overlooked on the premises that they do not directly contribute to recruitment and therefore to population growth rate, even in monogamous species (Caizergues & Ellison, 1997, Johnson, 2009, Schamber et al., 2009, Wilson & Martin, 2012, Gerber & White, 2014, Nicol-Harper et al., 2023, Soriano-Redondo et al., 2023, Viana et al., 2023). Yet changing sex structure is a reliable proxy of population dynamics/viability (Alexander & Taylor, 1983, Folliot et al., 2020), as well as a key driver of parenting and an evolutionary force of life history traits (Eberhart-Phillips et al., 2018, Székely & Miranda, 2026). Secondly, another implicit albeit unassessed assumption, is that temporal changes in sex structure can result from sex-specific changes of parameters such as survival or alteration of sex ratio at birth only (see Fox et al., 2016, Eberhart-Phillips et al., 2017, Ramula et al., 2018, Payevsky, 2021, Wood et al., 2021).

Common pochard (hereafter CP) populations of the North east/North west Mediterranean flyway have been declining since the end of the last century (Schricke, 2002). Recent studies not only confirmed a 4.9% short term annual decline in this flyway (Folliot et al., 2022), but also that an increase in proportion of males has occurred during the same period, rising from 62% to 71% in the mid-2010 (Carbone & Owen, 1995, Brides et al., 2017, Pöysä et al., 2019). European duck experts hypothesized that increasing mortality of incubating females and nests, owing to increasing predation pressure, likely explained the CP’s decline and the increasing proportion of males together (Fox et al., 2016). Recently, however, an analysis of capture-recapture-recoveries from Great-Britain, Switzerland and France, partly falsified this hypothesis, failing to detect any mortality increase over the period of decline (Folliot et al., 2020). Looking for an alternative hypothesis, the experts thus proposed that extensive water/sediments contamination by endocrine disruptors chemicals might have the potential to disrupt hormone deposits in eggs (Jouanneau et al., 2023) and alter gene expression (Brand et al., 2025), potentially causing a decrease in survival of female embryos (Tartu et al., 2014) that could explain both the decline of the species and the increasing proportion of males (Arzel et al., 2024).

In this paper, we use the case study of the declining populations of CP in Northwest Europe to emphasise the importance of both, prospective analyses and demographic models accounting for sex structure, for assessing the underlying mechanisms of resilience/collapse of populations facing multiple anthropogenic stressors. We compiled estimates of demographic parameters (survival, clutch size, nest survival…) of CP populations of the North east/North west Mediterranean flyway to parameterise a two-stage, two-sex population model to identify the drivers of demography and simulate both the trajectories of populations and variations of sex ratio under different scenarios (Usher, 1971, Caswell, 2019). As some key demographic parameters were not available, and most the others could not be considered as fully representative of the focal population (see material and methods), we could not perform a reliable population diagnostic. Our aim, therefore, was to build a model able to “enlighten” the properties of a CP population for identifying the parameters driving demography and explaining the decline.

More specifically, the scope of our study was threefold: 1) to assess the hypotheses put forward by duck experts for explaining the decline of CP (Fox et al., 2016) in anticipation of the implementation of adaptive harvest management (Månsson et al., 2023, Guillemain & Salas, 2024), 2) to emphasize the importance of considering males and females separately in monitoring schemes, and then the necessity to explore the conditions underlying variations of sex ratio in demographic models, and, 3) to provide recipes and derive generalities from our findings, for example, for evaluating the current status of other species for which data about changes in sex ratio are currently available or could be easily gathered (see e.g., Wood et al., 2021).

## Material and methods

### Origins and computation of demographic parameters

We compiled estimates of survival parameters of the flyway available in the literature and used reproductive parameters estimated from our own monitoring schemes in France (for details on field and statistical methods, see Bourdais et al., 2015, Folliot et al., 2017). As in many other duck species, juvenile CPs do not achieve a full moult until their first year of life and hence can be distinguished from 2 years-old or older individuals before reaching 14 months, by the shape and pattern of the primary wings feather (Mouronval, 2016). Therefore, following Blums et al. (1996), we assigned individuals to three categories: hatch year or HY aged of less than 10-11 months; second year individuals 10-11 to 22-months-old hereafter called SY, and, individuals ≥ 2 years-old or ASY, for after second year.

HY survival (survival from hatching to the breeding season of the next year) data are not only unavailable for CP, but have also rarely been estimated in other diving ducks (appendices). Therefore, we computed estimates of HY survival by combining the few estimates available in the literature for Lesser scaup *Aythya affinis* ducklings (Dawson & Clark, 1996) and winter survival of Common pochard HYs (Gourlay-Larour et al., 2014, Herrera, 2019), and obtained 2 values, 0.15 and 0.17 (see appendices). We discarded HY survival probabilities estimated in Blums et al. (1996) because they proved incompatible with the observed population dynamics of the population of Engure (appendices). Concerning this dataset, only CV was kept. SY survival was derived from Blums et al. (1996) and also computed using seasonal estimates of monthly survival in Gourlay-Larour et al. (2014) as 0.66 (appendices, Table 1). ASY survival was derived from capture-mark-recapture/recoveries studies conducted on individuals ringed in Northwest Europe, i.e., Switzerland ‘CH’, Great Britain ‘UK’ and France ‘FR’, (Folliot et al., 2020), and Latvia ‘LV’ (Blums et al., 1996) (Table 1).It should be noted, that all the survival estimates used in our study were computed from individuals belonging to harvested populations (see, Hirschfeld et al., 2019, Folliot et al., 2020), they therefore include the potential partially additive effects of hunting mortality (see Grzegorczyk et al., 2024).

**Table 1.** Survival parameters estimates of Common pochards by age (0 to ca 10-months-old = first year or HY, 1 to 2 to years-old = second year of SY, ≥ 2-years-old = after second year or ASY) and sex of Common pochards in Northwest Europe. (*) Value estimated on females extrapolated to males. (**) aberrant value not used for computed the average used as parameter in our models. (***) includes Engure’s **CV**.

| Age | Sex | Value | Conf. Int. | Range / Coefficient of process variation (CV) | Site | Time windows | References |
| --- | --- | --- | --- | --- | --- | --- | --- |
| HY | Males | 0.15 | - | Time & Sex independent | Dombes, France / North-America | - | Herrera 2019 / Dawson & Clarke 1996 |
|  |  | 0.17 | - | Time & Sex independent | Grand-lieu, France / North-America | - | Dawson & Clarke 1996 / Gourlay-Larour et al 2014 |
|  |  | 0.06 (**) | - | 0.03-0.14 / 0.51 | Engure, Latvia | 1976-1991 | Blums et al. 1996 |
|  |  | **0.16** |  | **0.03-0.17 / 0.26 (\*\*\*)** | **Average/Global** |  |  |
|  | Females | 0.15 | - | Time & Sex independent | Dombes, France / North-America | - | Herrera 2019 / Dawson & Clarke 1996 |
|  |  | 0.17 | - | Time & Sex independent | Grand-lieu, France / North-America | - | Dawson & Clarke 1996 / Gourlay-Larour et al 2014 |
|  |  | 0.06 (**) | - | 0.03-0.14 / 0.51 | Engure, Latvia | 1976-1991 | Blums et al. 1996 |
|  |  | **0.16** |  | **0.03-0.17 / 0.26 (\*\*\*)** | **Average/Global** |  |  |
| SY | Males | 0.55 (*) | - | 0.46 - 0.66 / 0.12 | Engure, Latvia | 1976-1990 | Blums et al. 1996 |
|  |  | 0.59 (*) | - |  | Engure, Latvia | 1976-1990 | " |
|  |  | 0.66 | 0.61-0.74 | Time & Sex independent | Grand-lieu, France | 2004-2011 | Gourlay-larour et al. 2014 |
|  |  | **0.60** |  | **0.46 - 0.66 / 0.13** | **Average/Global** |  |  |
|  | Females | 0.55 | - | 0.46 - 0.66 / 0.12 | Engure, Latvia | 1976-1990 | Blums et al. 1996 |
|  |  | 0.59 |  |  | Engure, Latvia | 1976-1990 | " |
|  |  | 0.66 | 0.61-0.74 | Time & Sex independent | Grand-lieu, France | 2004-2011 | Gourlay-larour et al. 2014 |
|  |  | **0.60** | - | **0.46 - 0.66 / 0.13** | **Average/Global** |  |  |
| ASY | Males | 0.66 | - | 0.52-0.80 / 0.13 | Grand-lieu, France | 2004-2017 | Folliot et al. 2020 |
|  |  | 0.75 | 0.74-0.76 | Time independent | UK | 1977-2011 | " |
|  |  | 0.81 | 0.80-0.82 | Time independent | CH | 1977-2011 | " |
|  |  | **0.74** | - | **0.52 - 0.81 / 0.12** | **Average/Global** |  |  |
|  | Females | 0.62 | - | 0.50-0.76 / 0.14 | Grand-lieu, France | 2004-2017 | Folliot et al. 2020 |
|  |  | 0.69 | 0.66-0.72 | Time independent | UK | 1977-2011 | " |
|  |  | 0.67 | 0.64-0.70 | Time independent | CH | 1977-2011 | " |
|  |  | 0.65 | - | 0.54 - 0.78 / 0.12 | Engure, Latvia | 1976-1990 | Blums et al. 1996 |
|  |  | 0.59 | - | - | Engure, Latvia | 1976-1990 | " |
|  |  | **0.644** |  | **0.50 - 0.78 / 0.12** | **Average/Global** |  |  |

Breeding probability and sex ratio at hatching are derived from Blums et al. (1996) and Blums and Mednis (1996), respectively (Table 2). We used the reproductive parameters (clutch size **CS**, nest survival **NS**) recorded: 1) on Grand-lieu lake, France, 2008-2017 and re-analysed for the purpose of this article (Folliot et al., 2017, Herrera, 2019), and, 2) in the Sologne fishponds area, France, 2014-2016 (Bourdais et al., 2015). We also included the estimate of nest survival on Engure lake, LV provided in Blums et al. (Blums et al., 1997) (see Table 2). Concerning the Grand-lieu dataset, which was the only site providing age-related estimates of **CS** and **NS**, only the samples (years) including at least 7 SYs were kept for computing average estimates and their respective coefficient of variation, that is 2006-2008 and 2010-2014 (Table 2).

**Table 2.** Breeding parameters of Common pochard females in the Northwest Europe by age (1-year-old = SY, ≥ 2-years-old = ASY). **PB** for probability of breeding, **sr** for sex ratio (here females/(males+females)), **CS** for clutch size, and **NS** for nest survival. Data on female embryos survival Se currently unavailable.

|  | Age | Value | Range /<br>Coefficient of process<br>variation | Site | Time window | References |
| --- | --- | --- | --- | --- | --- | --- |
| **PB** | SY | 0.7 | - | Engure, Latvia | 1976-1992 | Blums et al. 1996 |
|  | ASY | 1 | - | Engure, Latvia | 1976-1992 | " |
| **sr** |  | 0.498 | - | - | 1978-1993 | Blums & Mednis 1996 |
|  |  |  |  | - | - |  |
| **CS** | SY | 8.3 | 7.5-10.1 / 0.11 | Grand-lieu, France | 2006-2007 & 2010-2014 | Present study |
|  | ASY | 9.3 | 8.7-10.2 / 0.08 | Grand-lieu, France | 2006-2007 & 2010-2014 | Present study |
| **NS** | SY | 0.33 | 0.15-0.69 / 0.55 | Grand-lieu, France | 2006-2007 & 2010-2014 | Herrera 2019 |
|  | ASY | 0.48 | 0.30-0.79 / 0.33 | Grand-lieu, France | 2006-2007 & 2010-2014 | Herrera 2019 |
|  | SY+ASY | 0.32 | 0.18-0.44 / 0.41 | Sologne, France | 2012-2014 | Bourdais et al. 2016 |
|  | SY+ASY | 0.95 | Time & age independent | Engure, Latvia | 1978-1993 | Blums et al 1997 |
|  | **Average SY** | 0.53 | 0.15-0.95 / 0.45 |  |  |  |
|  | **Average ASY** | 0.58 | 0.30-0.95 / 0.35 |  |  |  |

Table 1 and 2 display the average estimates by site and age class, as well as the average values over sites and time (spatio-temporal averages), which were used to parameterise our demographic model. Whenever possible, i.e. when the parameter was found to vary across sites or time (years), we computed its coefficient of process variation (standard deviation to mean ratio of the process variation) which was then used in to compute true elasticities. Thus, our estimates of coefficient of process variation (**CV**) combine temporal and spatial environmental stochasticity (see also Table 3).

**Table 3.**
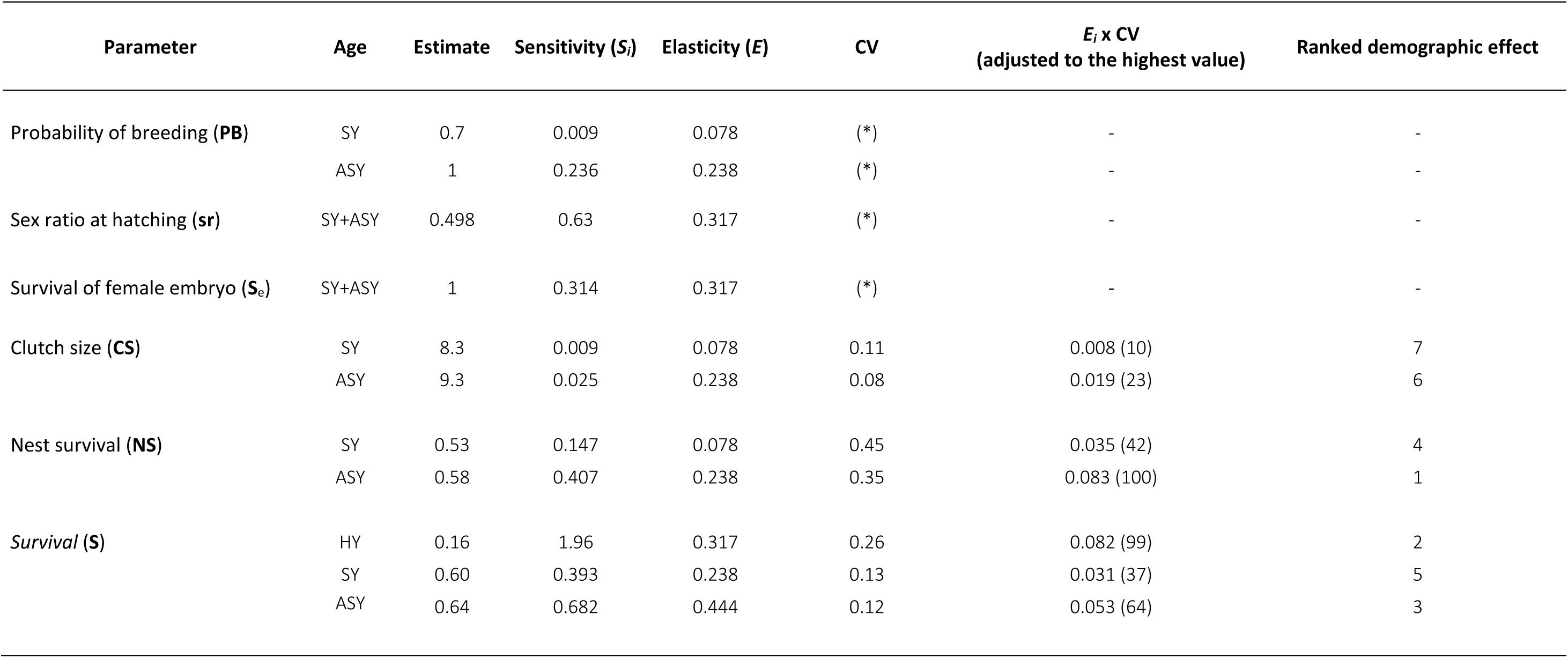
Sensitivities, elasticities of ***λ***, to variations in demographic parameters, together with elasticities pondered by the coefficient of process variation (**CV**) of parameters or true elasticities. The higher true elasticity, the greater the variations in ***λ*** are influenced by the parameter. Rank of demographic effects are derived from elasticity ***Ei*** x **CV** scores. (*) **CV** unknown

| Parameter | Age | Estimate | Sensitivity ($S_i$) | Elasticity ($E$) | CV | $E_i \times CV$<br>(adjusted to the highest value) | Ranked demographic effect |
| --- | --- | --- | --- | --- | --- | --- | --- |
| Probability of breeding (**PB**) | SY | 0.7 | 0.009 | 0.078 | (*) | - | - |
|  | ASY | 1 | 0.236 | 0.238 | (*) | - | - |
| Sex ratio at hatching (**sr**) | SY+ASY | 0.498 | 0.63 | 0.317 | (*) | - | - |
| Survival of female embryo (**S<sub>e</sub>**) | SY+ASY | 1 | 0.314 | 0.317 | (*) | - | - |
| Clutch size (**CS**) | SY | 8.3 | 0.009 | 0.078 | 0.11 | 0.008 (10) | 7 |
|  | ASY | 9.3 | 0.025 | 0.238 | 0.08 | 0.019 (23) | 6 |
| Nest survival (**NS**) | SY | 0.53 | 0.147 | 0.078 | 0.45 | 0.035 (42) | 4 |
|  | ASY | 0.58 | 0.407 | 0.238 | 0.35 | 0.083 (100) | 1 |
| Survival (**S**) | HY | 0.16 | 1.96 | 0.317 | 0.26 | 0.082 (99) | 2 |
|  | SY | 0.60 | 0.393 | 0.238 | 0.13 | 0.031 (37) | 5 |
|  | ASY | 0.64 | 0.682 | 0.444 | 0.12 | 0.053 (64) | 3 |

### Demographic model and parameters contribution to population growth rate

Our objective to provide −given available data− the most detailed and robust demographic model for extracting reliable demographic properties (generation time, sensitivities ***S_i_*** and elasticities ***E_i_*** of ***λ*** to change in a particular demographic parameter or set of parameters) of a typical Common pochard population. Thanks to a quite detailed set of data available (Tables 1 and 2), we could build a life-cycle that included both sexes as well as 2 stages (ages classes, Figure 1).

**Figure 1.**
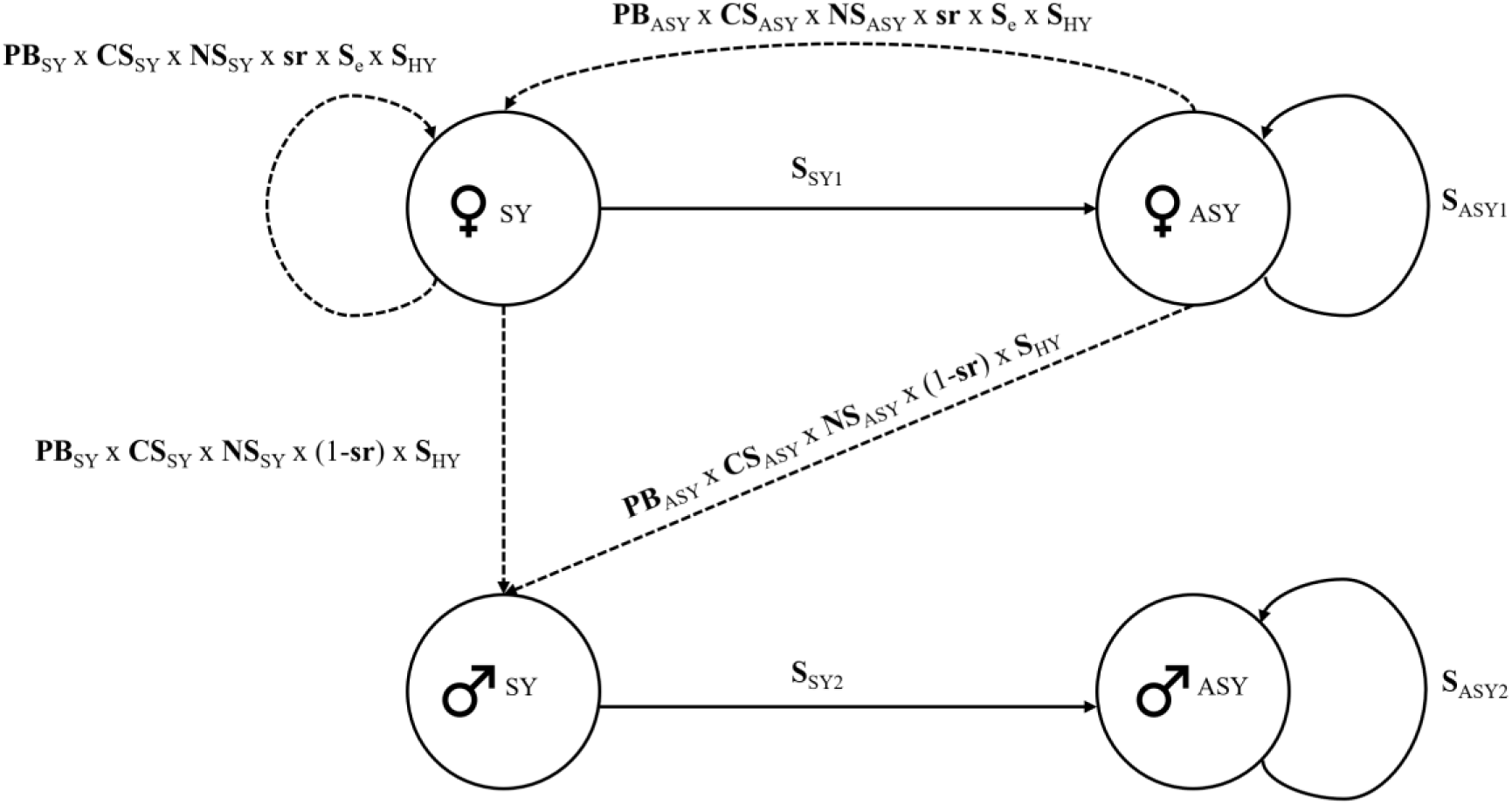
The life cycle of the deterministic Leslie-Usher Matrix model (**A**) used in the present study. Subscript **1** and subscript **2** are for females and male respectively (the absence of subscript means that the parameter takes the same valuer for males and females). HY is for 0 to 10-11 months individuals, SY for 10-11 to 20-22 months individuals, ASY for ≥ 2-years-old individuals, and e stand for female embryos (see **S**e embryo female survival). The parameter **S** is for survival probability, **PB** for probability of breeding, **CS** for clutch-size, **sr** for sex ratio at hatching and **NS** for nest survival. Parameters used in the model are provided in Tables 1 and 2. **S**e initial value’s set at 1.

We implemented this lifecycle using a before reproduction Leslie-Usher projection matrix **A** with a 1year time step (Leslie, 1945, Usher, 1971, Caswell, 2000, Caswell, 2019).

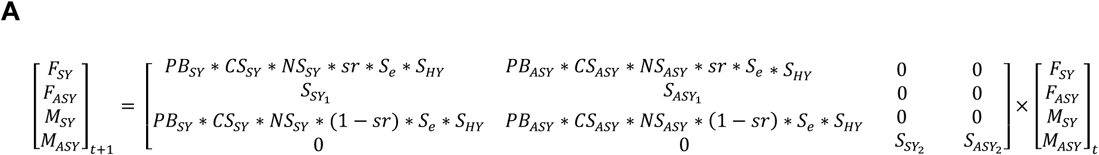

For the sake of simplicity, we opted for a deterministic analysis of sensitivity of ***λ*** to changes in the demographic parameters. We first computed the effects of absolute changes in each demographic parameter on ***λ*** (sensitivities, ***S_i_***), then their relative effects (elasticities, ***E_i_***) (Caswell, 2000, Caswell, 2019). We used the product of the elasticities of parameters with their respective coefficient of process variation to rank each parameter according to its potential driving effects on population growth rate ***λ*** (Steen & Erikstad, 1996). Although this last approach is effective for identifying the factor(s) driving ***λ***, assessing the potential demographic effect of parameters may be best achieved by performing a disturbance analysis thereby the impact of each parameter on ***λ*** is assess by making it vary when the over ones are held constant. Indeed, some parameters are upper-bounded (adult survival), while others are less so (HY survival), which let more room for them to increase and hence to be acted on.

We used the R package ‘popbio’ (R Core Team, 2023) to implement the projection matrix and to extract its various properties including the highest order eigen value which is the population growth rate ***λ*** with lambda() and to derive sensitivities, and first and second order elasticities with the vitalsens() function. Elasticities of first order parameters sum to 1 and give the relative contribution of these parameters to ***λ***. Elasticities to second order parameters do not sum to 1 and show how much a proportional change in any given such parameter affect ***λ*** (see section 9 in Caswell, 2000). We extracted ***λ*** under various combinations of parameters to compare graphically the relative influence of key parameters.

### Combined effects of recruitment components to *λ* and male proportions

One aim of our paper was to evaluate the various hypotheses put forward by experts for explaining the decline of CP in North east/North west Europe (Fox et al., 2016, Brides et al., 2017, Folliot et al., 2020). One of these hypotheses −suggesting that declining survival of mature females during breeding owing for example to increased predation pressure would be involved− has already been discarded by Folliot et al. (2020) and will be therefore not explored in details here. According to available data, the decline of CP in Northwest Europe was accompanied by an increase in male proportions from ca 0.61 to ca 0.71 (Brides et al., 2017). Thus, any change in any demographic parameter or combination of parameters had to explain the population decline and increasing proportions of males together. This leaved us with two possibilities: 1) decreasing proportions of females among hatchlings (secondary sex ratio), e.g., due to decreasing survival of female embryos **S**_e_ (Arzel et al., 2024), or eventually to changes in primary sex ratio due to genetic females developing into phenotypic males, 2) decreasing nest survival and/or duckling survival as hypothesised by Folliot et al. (2020). We assessed these hypotheses by simulating the effect of changes in these candidate parameters on both ***λ*** and male proportions extracted from the projection matrix **A**.

## Results

### Contribution to population growth rate (demographic drivers)

Using the average demographic parameters in Tables 1 and 2, we obtained an estimate of population growth rate very close to stability (***λ*** = 0.99). The mean generation length ***T*** (average age of females) of 3.12 years. The elasticity analysis showed: 1) that survival of SY and ASY taken together contributed to a larger part of ***λ*** (68.3%) than their components of recruitment (**CS**, **NS** and **S**_HY_, 31.7%), and 2) that ASY contributed to a larger part of ***λ*** (68.3%) than SY (31.7%, see Table 3). However, the driver effects of productivity parameters on the demography were not negligible when environmental stochasticity (coefficient of process variation **CV**) was accounted for. For example, although the sum of **NS** elasticities of SYs and ASYs place this parameter at the second rank in terms of impact on ***λ*** (0.32 against 0.68 for the combined elasticities of survival probabilities of SYs and ASYs), it is in fact most potent than adult survival as driver of population dynamics when **CV** is accounted for (***E*_NS(_**_sy asy combined)_ **x CV_NS_**_(sy asy combined)_ = 0.118 against 0.081 for ***E*_S_**_(sy asy combined)_ **x CV_S_**_(sy asy combined)_, see Table 3). Among all demographic parameters included in our transition matrix, **CS** has the less impact on ***λ***, because it displays both a modest elasticity and a low **CV** at the same time (Table 3). PB, **sr** and **S**_e_ could be potent drivers of ***λ***, but their process variances have not been estimated so far (see next section).

Overall, the perturbation experiments (Figure 1) confirm the elasticity analysis (including that **NS** is the driver and limiting parameter). This time however, first year survival **S**_HY_ clearly appears to have the highest potential impact on ***λ*** being almost twice that of combined survival of SYs and ASYs (Figure 1). Thus, if any management policy could improve first year survival, even modestly, it would trigger the greatest demographic response. Both, **S**_e_ and especial sex ratio (female/male) at hatching (due to female masculinisation or male feminisation at the embryo stage) have a potential strong influence (although much weaker than **S**_HY_, see Figure 1), but their potential of variation under anthropogenic pressure, such as EDC, is unknown. Finally, increasing PB of SY females –assuming it is possible– would have a very limited impact on ***λ*.**

**Figure 1.**
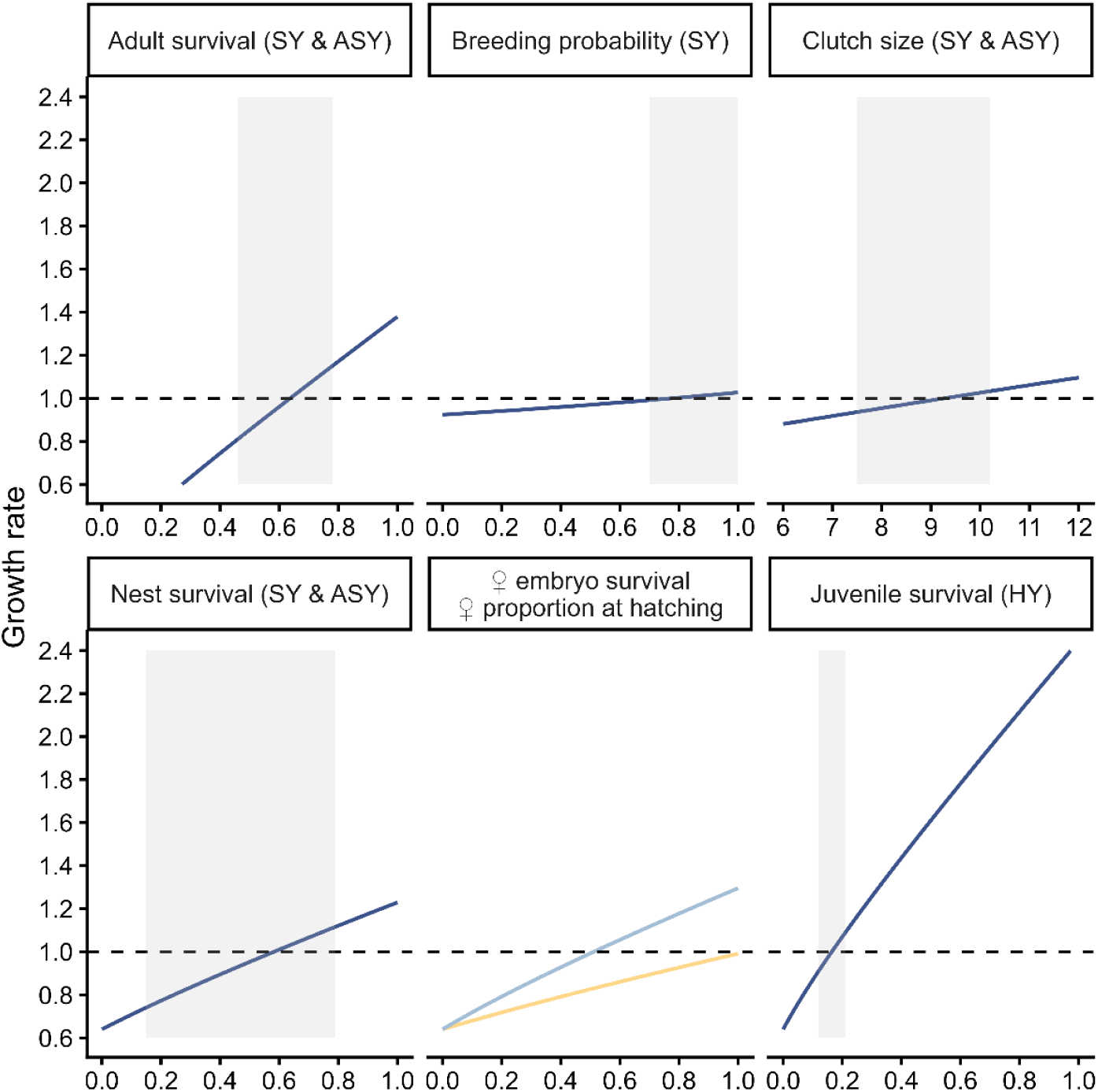
Effects of survival, probability of breeding, clutch size, nest survival of matures (SY and ASY) Common pochard females and sex ratio at hatching, survival of female embryos and survival of hatch year females (HY), on population growth rate ***λ***. The grey shaded areas show the annual range of variation of the parameter (due to environmental stochasticity). The dashed line represents the demographic equilibrium (***λ*** *= 1*).

### Combined effects on *λ* and sex structure

The proportion of males derived from our projection matrix is male biased (58%). Decreasing **NS**, **S**_HY_ or **S**_e_ have the potential to affect both ***λ*** and the proportions of males among breeders (***P*_♂_**) to quite large extent (Figures 1 & 2). However, according to our simulations a drop of population growth rate of 4-5%, like the one derived from counts at the level of the flyway –obtained by decreasing reproductive success of 0.1 in both SYs and ASYs or **S**_HY_ by 0.02– yield a proportion of males of 59% which is much lower than the estimate of Brides et al. (2017) of 71%. Similarly, decreasing proportions of females at hatching **sr** (due to genetic female embryos developing into phenotypic males) have a strong effect on both of these parameters (Figure 1 & 2), eventually matching more or less the values derived from counts. For example, for **sr** = 0.4, the derived values of ***λ*** and ***P*_♂_** were 0.94 and 0.69 respectively, that is, very close to the estimates of 0.95 and 0.71 of Folliot et al. (2022) and Brides et al. (2017), respectively.

Nonetheless, one major finding of these simulations worth emphasizing –both because not intuitive and with important consequences for assessing population status– is that parameters indiscriminately affecting productivity of individuals of both sexes (**NS** and **S**_HY_) can have an impact on ***P*_♂_** almost as high as sex-specific parameters such as **sr** and **S**_e_ (see Figure 3). This is possible because adult males experience much higher survival probability than females (0.74 against 0.64). Thus, when productivity/recruitment decreases (owing to decreasing **NS** or **S**_HY_), the weight of adults on the sex structure of the population “mechanically” increases, giving “larger room” for the between-sex difference in adult survival to express itself and to affect ***P*_♂_**. It ensues, that low levels of one of the over of the components of recruitment (especially **NS** and **F**_HY_) can lead to highly male-biased sex-ratio (Figure 2).

**Figure 2.**
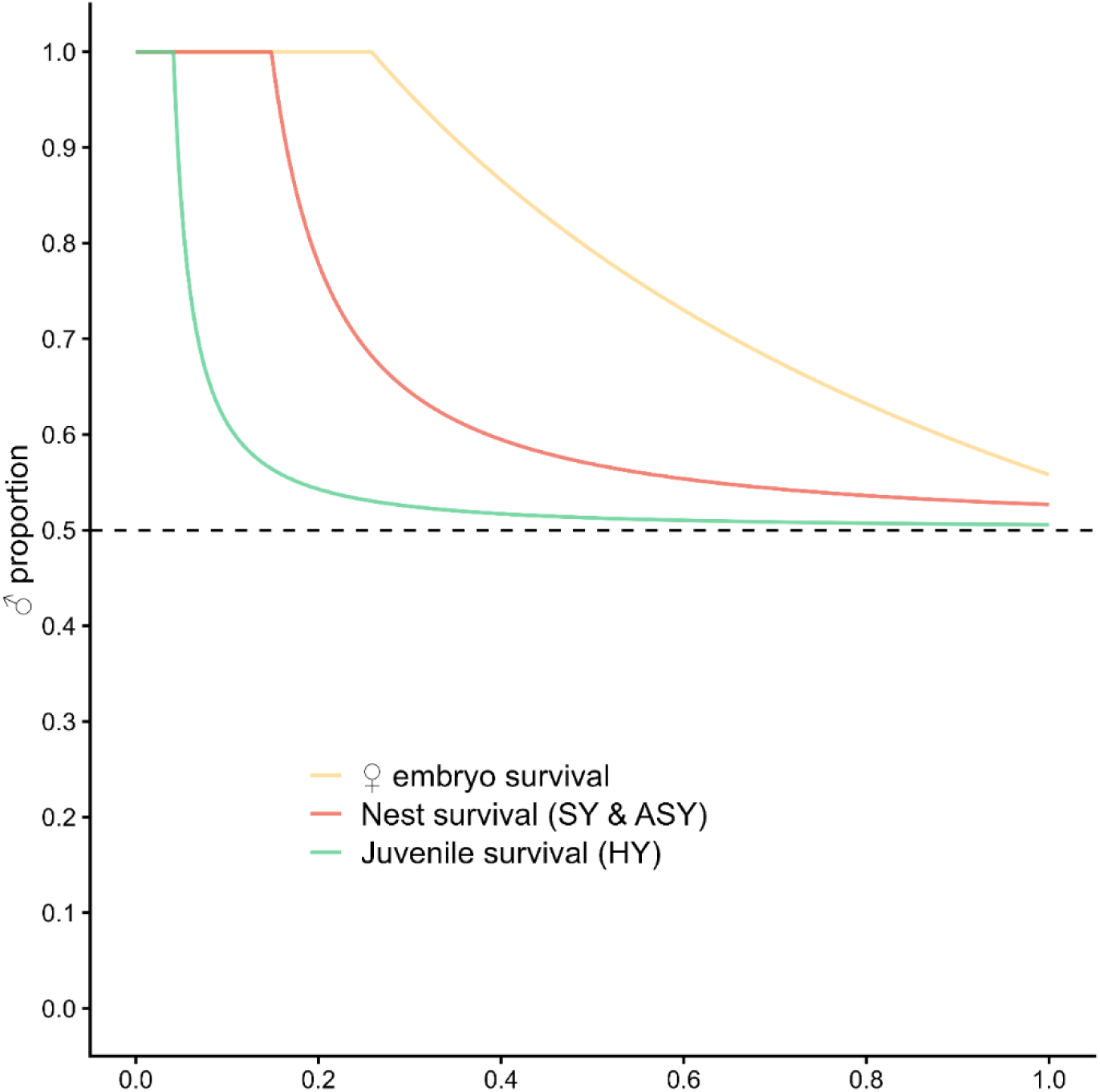
Effects of changing of three components of recruitment (**S**e, **S**HY, **NS**) that have been hypothesized to potentially account for the decline of Common pochard populations in North east/North west Europe, on the proportions of males among adults. Interestingly, because among adults, male survival is higher than female survival, proportions of males can increase even if when both sexes suffer the same decline in recruitment (ie., when **NS** or **S**_HY_) decrease).

**Figure 3.**
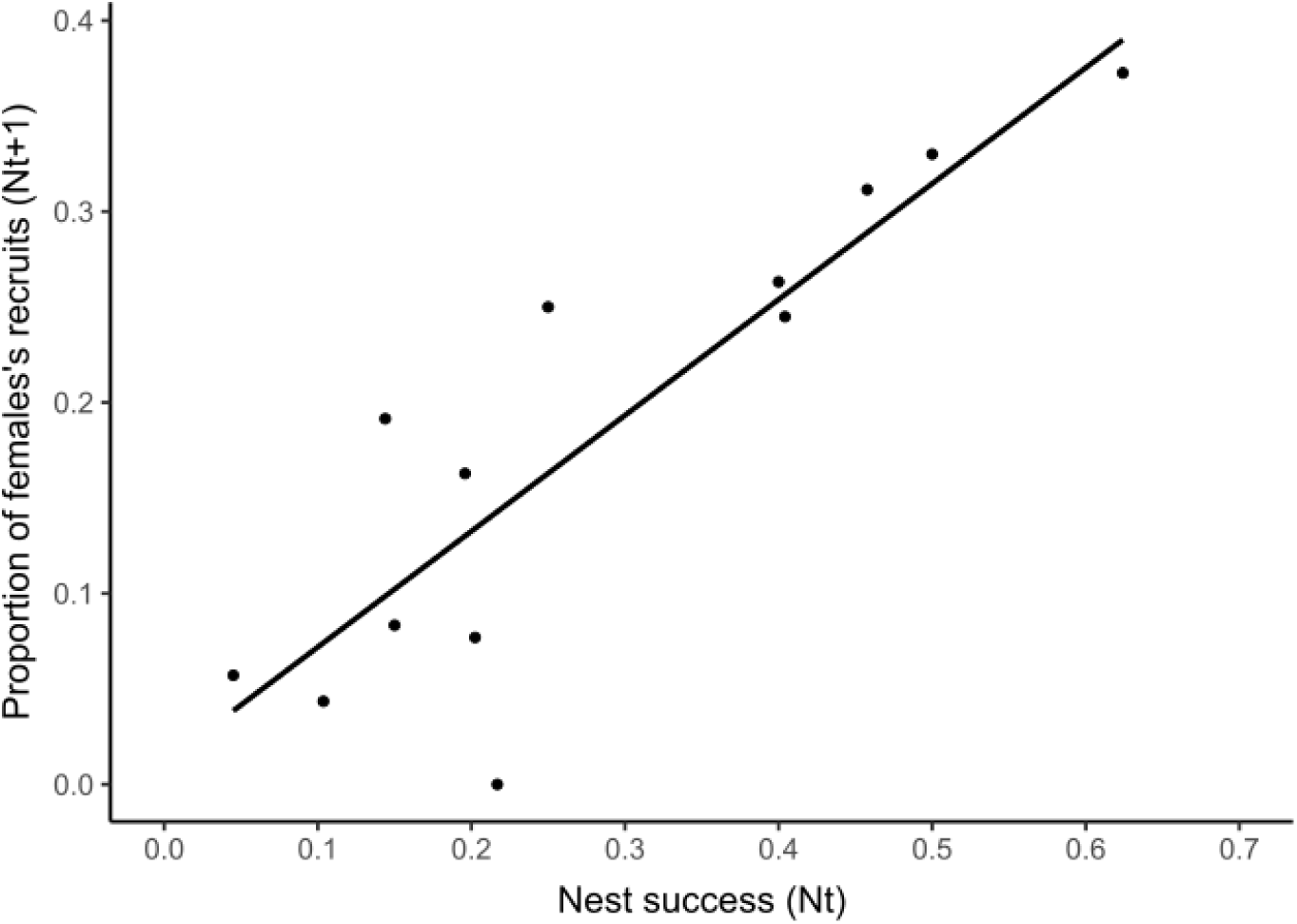
On Grand-lieu lake (France) there was a relationship between nest survival in year N and the proportions of yearling females among nesting females in year N+1, supporting that nest survival was an important driver of CP populations.

## Discussion

The present study provides key insights into the understanding of the mechanisms/drivers underlying the demography of CP, the identification of potential causes of decline and, more importantly, the necessity of accounting for male in demographic studies.

### Population drivers, cause of the decline and management options

With a generation time slightly above 3 years, the Common pochard is rather on the slow side of the slow-fast continuum (see e.g., Lebreton & Clobert, 1991, Koons et al., 2014, Flint, 2015). This was confirmed by the elasticity analysis showing that combined survival of SYs and ASYs contributed to almost 70% of the population growth rate. However, graphic representations of perturbation analyses clearly pinpoint first year (HY) survival as key component of ***λ*** (Figure 1). Altogether therefore, our study support that improving survival would be a particularly effective way to increase ***λ*** of CP populations.

However, being more sensitive to changes in survival does not necessarily implies that this parameter drives demography (Koons et al., 2014, Flint, 2015, Hepp et al., 2020). On the contrary, according to the demographic buffering hypothesis, slow species, whose demography is primarily sensitive to changes in survival of mature females, tend to buffer survival from challenging environmental conditions at the expense of the component of recruitment (nesting success, juvenile survival…, see Pfister, 1998, Gaillard & Yoccoz, 2003, Rotella et al., 2012, Hilde et al., 2020). In such species, trading current reproduction against higher future reproductive prospects would pay more in terms of fitness than the reverse strategy (Trivers, 1974, Hilde et al., 2020). In this respect, the CP is not exception to the rule. Indeed, despite ***λ*** displays a weaker elasticity to nest survival than to survival of mature females, the first experiences a much larger environmental stochasticity, making it the primary driver of changes in numbers of CP. This finding is supported by strong empirical evidence showing that nest survival does affect recruitment (Figure 3 below). Altogether our data, therefore support that declining nest survival is an important driver of CP demography and thus could be involved in the decline observed in North east/North west Europe.

Our study suggests that **S**_HY_ is another critical driver of CP’s populations. Unfortunately, reliable estimates of **S**_HY_ and its process variance are lacking. Examples of the importance of components of recruitment (such as **NS** and **S**_HY_) as demographic drivers of species with high adult survival, have been gathered in numerous medium-sized birds including Tetraonids (Caizergues & Ellison, 1997, Rotelli et al., 2021, Canonne et al., 2023) and ducks (Koons et al., 2014, Flint, 2015, Arnold et al., 2018, Hepp et al., 2020, Wiegers et al., 2022, Gibson et al., 2025).

Hunting is a major cause of mortality of ducks, including CP (Guillemain et al., 2016, Hirschfeld et al., 2019, Souchay et al., 2026). Moreover, hunting mortality is probably partly additive to natural mortality (Grzegorczyk et al., 2024, Souchay et al., 2026), especially in adult females supposed to suffer higher losses during breeding (Krementz et al., 1987, Arnold et al., 2012, Gibson et al., 2025) and first year individuals which are particularly vulnerable to hunting (Gibson et al., 2025, Souchay et al., 2026). Thus, in the short term, hunting ban could be a more effective and more tractable way to improve population growth rate than targeting nest survival (see Flint, 2015), even if decreasing nest survival is the most important component of the decline. Examples of hunting ban of females include lekking grouses (Black grouse and Capercaillie), whose females benefit from a status of total protection (Caizergues & Ellison, 1997) and the Common Eider, for which the hunting ban of female has been recently implemented in numerous countries (see Tjørnløv et al., 2019).

### Changing proportions of males in sexually matures individuals does imply sex-specific changes in survival at some stage of the life-cycle

Adult sex ratio is usually male-biased in ducks (Owen & Dix, 1986, Carbone & Owen, 1995, Pöysä et al., 2019, Gibson et al., 2025) and, following the hypothesis initially put forward by Bellrose et al. (1961), it is now widely accepted that the excess of males among adults results from the lower survival of females during the breeding season (Conroy & Eberhardt, 1983, Krementz et al., 1987, Brasher et al., 2006, Lehikoinen et al., 2008, Arnold et al., 2012, Bellebaum & Mädlow, 2015, Pöysä et al., 2019, Gibson et al., 2025). Indeed, there is ample empirical evidence that males experience higher survival than females in both diving and dabbling ducks (e.g., Lake et al., 2006, Reynolds & Citta, 2007, Gunnarsson et al., 2008, Devineau et al., 2010, Sun et al., 2011, Milton et al., 2016, Bielefeld et al., 2020, Folliot et al., 2020). The causes of male-biased sex ratio are therefore well established in many ducks, and it has become implicit that changing sex ratio could only result, either from an alteration in primary sex ratio, or from changes in sex-specific survival at some stage of the life cycle (e.g., Lehikoinen et al., 2008, Fox et al., 2016, Ramula et al., 2018, Ellis et al., 2022). Yet, Folliot et al. (2020) showed that changes in nest survival –a parameter affecting equally male and female recruitment– could also alter adult sex ratio (see also Alexander & Taylor, 1983). Our simulations confirm this view, providing convincing evidence that: 1) the excess of males results from a lower survival of females among sexually matures individuals and, more importantly and, 2) as a consequence, *any change in any factor affecting recruitment of both sexes equally* (nest survival, post-fledging survival…) *is sufficient to induce both, a decline in **λ** and an increase of male proportions among sexually matures individuals*. In other words, *there is no need to call for any change in sex-specific survival at different life stages for triggering/explaining changing sex ratio when between-sex differences of survival are observed among adults*. More specifically, when among adults, male survival is higher than female survival, proportions of males increase when recruitment decreases and reciprocally. The reverse would be true when adult females survive better than males: decreasing recruitment would trigger increasing proportions of females. The corollary of this finding, is that in species like ducks, characterised by higher survival of males than females among adults, increasing proportions of males should be, by default, interpreted as indicators of recruitment problems, particularly when they are accompanied with populations’ declines. There is a scientific consensus that components of (female) recruitment –more particularly nest survival and first year survival– are potent drivers of ducks’ population dynamics (Hario et al., 2009, Schamber et al., 2009, Rönkä et al., 2011, Öst et al., 2016, Koons et al., 2017, Arnold et al., 2018, Hepp et al., 2020, Tjørnløv et al., 2020). Moreover, in support to our finding, there is not only empirical evidence that proportions of males have recently increased in many duck species, including CPs (e.g., Christensen & Fox, 2014, Brides et al., 2017, Wood et al., 2021, Warrender et al., 2026), but also that these increases most probably result from decreasing recruitment (Christensen & Fox, 2014). This view is further supported by the fact that in neither of these studies was any bias (or any change) of sex ratio detected among first year individuals (Christensen & Fox, 2014, Wood et al., 2021).

Of course, our simulations also show that any factor affecting sex ratio at hatching (e.g., increased mortality or masculinisation of female embryos) could greatly affect simultaneously the proportion of males among adults and population growth rates. However, although female masculinisation (e.g., in response to endocrine disruptors) is not implausible (Arnold & Itoh, 2011, Major & Smith, 2016) and sometimes observed under natural conditions (Chiba & Honma, 2011), neither its occurrence at the pre-hatching stage, nor its demographic impact have been investigated so far. Similarly, while in birds evidence exist about both, the transfer of endocrine disruptors from the mother to the egg, and the negative effect of mother’s contamination on hormone deposition into the egg (Jouanneau et al., 2023), little is known about their effects on sex-specific mortality of embryos (review in Ottinger et al., 2015). Thus, despite problems of masculinization and/or egg hatchability are sometimes reported in ducks, their potential causes and possible demographic consequences remain sketchy. Moreover, in diving ducks, the study of egg hatchability is blurred by the pervasive phenomenon of intraspecific nest parasitism, and problems of hatchability may befall primarily parasitic eggs (see Dugger et al., 1999, Šťovíček et al., 2013, Pöysä et al., 2014, Neužilová, 2015). Nevertheless, using microsatellites to distinguish parasitic from non-parasitic eggs Šťovíček et al. (2013), did not detected high egg mortality rates (i.e., 5%) among non-parasitic eggs in a CP population with very high rates of conspecific brood parasitism. Furthermore, under similar conditions, Blums & Mednis (1996) did not observe any departure from an even sex ratio. So far, therefore, empirical evidence of the possible impact of female masculinisation or female biased embryo mortality on the proportion of males among adult is at best circumstantial (see Arzel et al. 2024).

### Explaining the discrepancies between the model and observations

Among the candidate parameters assessed in our simulations, only sex ratio at hatching yielded a proportion of males among adults and a ***λ*** matching the estimates derived from counts by Brides et al. (2017) and Folliot et al. 2020 Folliot et al. (2020) respectively. However, the possible impact of sex ratio alterations (either through **sr** or **S**_e_) remains so speculative that the hypothesis can be discarded at least provisionally (see above). This leaves us with **NS** and **S**<u>_HY_</u>, those predicted values of male proportions are 0.59, when a drop of 4-5% of population growth rate is targeted while Brides et al. (2017) estimate was 0.71. The discrepancies in sex structure projected by our model and the value derived from counts could result from the fact that survival of SYs which were considered identical between sexes in our projection matrix (table 1) would be, in reality, male-biased too (e.g., Folliot et al., 2020). Alternatively, Brides et al. (2017), and Carbone & Owen (1995) before them, did not weight their local estimates of male proportions against local population size (a prerequisite for deriving a global value for the focal population) and therefore could have over-estimated male proportions. Thus, further works is needed, both, for assessing sex-specific survival parameters over the life-cycle and to provide unbiased sex ratio estimates from winter counts. Nevertheless, local proportions of males in a moderately declining population of Centre France (Sologne fishpond area), have recently been estimated at 62% (N=320, Caizergues et al. unpublished resultes), a value close to our own estimates under a 4.5-5% annual population decline.

## Conclusion and perspectives

Our study shows that populations of species sensitive to changing adult survival, may nevertheless be driven by nest survival because it usually experiences much higher environmental variance than adult survival (see e.g., Flint, 2015). However, just because nest survival is the driver of Common pochard populations doesn’t mean that management actions should target this parameter in priority. On the contrary, limiting mortality of females is potentially the most effective way to mitigate population declines of such species, first because of a much greater tractability (it is much easier to ban hunting than to improve habitat for nesting or brood rearing), and a much greater expected effect, especially concerning survival of juveniles after independence (due the much higher potential impact of first year survival on population growth rate). This is probably the reason why priority has almost always been given to hunting ban of females to restore populations of sexually dimorphic species such as lekking grouses (Caizergues & Ellison, 1997), or even ducks (Tjørnløv et al., 2019). However, this kind of rationale, entirely rests upon the assumption that hunting mortality limits both survival and population size. Hunting ban would be totally ineffective if CP populations’ decreases in productivity directly result from a decrease in the absolute number of nests (or young) that can be produced (carrying capacity is limiting productivity).

One important finding of this study concerns the importance to account for both sexes for unravelling the mechanisms underlying population dynamics and evolution (Gerber & White, 2014, Székely & Miranda, 2026). Increasing proportions of males observed in declining duck species, including the Common pochard, have become the focal point of many hypotheses all (explicitly or implicitly) assuming that changing sex structure (sex ratio) should result from sex-specific changes in demographic parameters only. Yet, our study clearly shows, that when sex-differences of survival exist among adults, changing recruitment (nest survival, first year survival…) affecting both sexes equally, can profoundly affect sex ratio too. In ducks, for example, increasing male proportions observed in many species (Christensen & Fox, 2014, Wood et al., 2021, Warrender et al., 2026) are likely the result of declining recruitment. Thus, assessing changing sex ratio in counts and accounting for males in demographic models are crucial steps towards a better understanding of the mechanisms underlying the dynamics an evolution of natural populations (Gerber & White, 2014, Székely & Miranda, 2026). In other words, although for different reasons, we concur with previous authors before us (Gerber & White, 2014, Székely & G. Miranda, 2026) that males should never be neglected in demographic or evolutionary studies as it is too often the case actually, and that monitoring schemes should strive harder at providing both (unbiased) sex ratio estimates, and sex-specific demographic parameters.

## Appendices

## After second year (ASY) survival estimates

Survival of individuals aged of 2 years or older (ASYs) are from (Blums et al., 1996) in Latvia (LV) and Folliot et al. (2020) in the United Kingdom (UK), Switzerland (CH) and France (FR). The LV dataset spanned over 16 years of ringing and physical recapture (1976-1990) on a single ringing site, the lake of Engure, as well as recoveries. The CH and UK datasets included 35 years of winter ringing (1977–2011) from a number of different ringing sites and dead recoveries only. The FR dataset consisted of 14 years (2004–2017) of winter ringing from a single ringing site (the lake of Grand-lieu, Northwest France) as well as recaptures and dead recoveries potentially anywhere within the Common pochard’s range. Terminology from Blums et al. (1996).

## Estimating first year (HY) and second year (SY) survival

Survival of ducklings from hatching to the moment when they enter the population the following year was taken as the product : 1) of survival of Lesser scaup (*Aythya affinis*) ducklings from 0 to 14 days-old (0.402, Dawson & Clark 1997) with the 10 months (August - May) survival of 1032 Common pochards ducklings ringed in France (Dombes fishponds) in the 1980-90s (0.37, Herrera 2019), that is ≈ 0.15, 2) of the same Dawson & Clark 1997 estimate (0.402) with their daily survival estimate of 15-48 days-old Lesser scaups (0.999) raised to the power of 62 days (to encompass daily survival over July-31 August) and overwinter monthly survival estimate of individuals ringed during their first winter of life Gourlay-Larour et al. (2014) raised to the power of 8 months (that is 0.402x0.999^62^x0.904^8^ ≈ 0.17.

Survival of SYs (from 10-12 months- to ≤ 2 years-old) was derived from SYs’ (15 May – 15 December) and ASYs’ (15 December – 15 May) monthly survival estimates in Gourlay-Larour et al. (2014) as 0.996^6^x0.937^6^=0.66.

Terminology from Blums et al. (1996).

## Assessing the validity of HY survival estimates in Blums et al. 1996

To assess the validity of parameters supposed poorly reliable, we ran preliminary analyses. More particularly, we ran a female only matrix model parameterised with the set of parameters of Engure, Latvia (excluding **CS** which was derived from Grand-lieu, France) with **S**_SY_ =0.55, **S**_ASY_ =**0.65**, **S**_HY_ = 0.06, **CS**_SY_ = 8.3, **CS**_ASY_ = 9.3, **NS**_SY_ = **NS**_ASY_ = 0.95, **PB**_SY_ = 0.7, **PB**_ASY_ = 1, **sr** = 0.4998, **S**_e_=1 (see Blums & Mednis, 1996, Blums et al., 1996, Blums et al., 1997), to assess the validity of the estimate of first year survival, that we suspected being underestimated compared to expectations (e.g., to values derived from by combining data of Dawson et al. 1997 and Hererra 2019, see Table 1). Using this set of parameters yielded a population growth rate ***λ*** = 0.85, incompatible with the trend of the local population of Engure (stable or increasing) which justified to discard the estimate of HY survival of Latvia from our analyses. Nevertheless, without any over alternative, we decided to keep CV of HY survival of Latvia as a proxy of CV of HY survival.

Blums P, Mednis A, Bauga I, Nichols J D, Hines J. E (1996) Age-specific survival and philopatry in three species of European ducks: a long-term study. *The Condor,* **98,** 61-74.
Dawson R D, Clark R G (1996) Effects of variation in egg size and hatching date on survival of lesser scaup (*Aythya affinis*) ducklings. Ibis 138:693-699.
Folliot B, Souchay G, Champagnon J, Guillemain M, Durham M, Hearn R, Hofer J, Laesser J, Sorin C, Caizergues A (2020) When survival matters: is decreasing survival underlying the decline of common pochard in western Europe? *Wildlife Biology,* **2020,** 1-12..
Gourlay-Larour M L, Pradel R, Guillemain M, Guitton J S, L’Hostis M, Santin-Janin H, Caizergues A (2014) Movement patterns in a partial migrant: a multi-event capture-recapture approach. *PLoS One*, ***9*** (5), e96478.
Herrera A. (2019) Estimations des taux de survies d’une population reproductrice de canards soumis à prélèvements: le cas du Fuligule milouin. Survival estimates of a breeding population of harvested ducks: the Common pochard as a case study. **Vol. MSc**, pp. 31. PSL Université Paris, Ecole Pratique des Hautes Etudes.

## Acknowledgements

This study has been possible thanks to the contribution of numerous anonymous ringers involved in the various ringing schemes in France and abroad. Special thanks to Christophe Sorin and the Fédération des Chasseurs de Loire-Atlantique for their invaluable help in Common pochards’ ringing on Grand-lieu lake. Many thanks to Serge Bourdais and the Fédération des Chasseurs du Loir-et-Cher for their involvement in nest monitoring.

## Data, scripts, code, and supplementary information availability

Data, scripts and supplementary information are available online (https://doi.org/10.5281/zenodo.19183773; Caizergues, et al. 2026)

